# Long-distance Wnt transport in axons highlights cell type-specific modes of Wnt transport *in vivo*

**DOI:** 10.1101/2023.05.03.539245

**Authors:** Ariel M Pani, Michelle Favichia, Bob Goldstein

## Abstract

Wnt signaling performs critical functions in development, homeostasis, and disease states. Wnt ligands are secreted signaling proteins that often move between cells to activate signaling across a range of distances and concentrations. In different animals and developmental contexts, Wnts utilize distinct mechanisms for intercellular transport including diffusion, cytonemes and exosomes [1]. Mechanisms for intercellular Wnt dispersal remain controversial in part due to technical challenges with visualizing endogenous Wnt proteins *in vivo*, which has limited our understanding of Wnt transport dynamics. As a result, the cell-biological bases for long-range Wnt dispersal remain unknown in most instances, and the extent to which differences in Wnt transport mechanisms vary by cell type, organism, and/or ligand remain uncertain. To investigate processes underlying long-range Wnt transport *in vivo*, we utilized *C. elegans* as an experimentally tractable model where it is possible to tag endogenous Wnts with fluorescent proteins without disrupting signaling [2]. Live imaging of two endogenously tagged Wnt homologs revealed a novel mode for long-distance Wnt movement in axon-like structures that may complement Wnt gradients generated by diffusion and highlighted cell type-specific Wnt transport processes *in vivo*.

---

A comprehensive understanding of Wnt dispersal mechanisms *in vivo* will require visualizing endogenous Wnt protein dynamics in living animals. To elucidate the processes involved in long-distance Wnt transport, we utilized spinning disk microscopy to characterize dynamics of two endogenously tagged Wnt homologs, Wnt/CWN-1::mNeonGreen (mNG) and Wnt/EGL-20::mNG in neurons during larval development. CWN-1 and EGL-20 are produced by multiple cell types in embryonic, larval, and adult animals including muscles and muscle progenitors, epithelial cells, and neurons [2, 3]. Previous studies of Wnt dispersal in *C. elegans* showed that endogenously tagged EGL-20::mNG forms a long-range protein gradient by extracellular diffusion during early larval development [2], and extracellular diffusion has also been reported for transgenically expressed CWN-1::YFP in embryos [4]. However, in examining Wnt/CWN-1::mNG and Wnt/EGL-20::mNG localization in later larval stages, we became intrigued by the possibility that neuronal cell architectures could provide routes for long-distance Wnt transport throughout the body independent of extracellular dispersal or specialized structures such as cytonemes.

Live imaging of endogenous Wnt/CWN-1::mNG and Wnt/EGL-20::mNG revealed two unexpected scenarios where Wnt proteins were delivered over long distances using distinct transport processes within axon-like structures or axons. We first observed that endogenously tagged CWN-1 is prominently expressed in Canal Associated Neuron (CAN) cells during larval development (Fig. 1a-c). Strikingly, CWN-1::mNG localized to mobile punctae within CAN axon-like structures (Fig. 1a-c, Videos S1, S2), which was distinct from localization in other Wnt-expressing cell types. Intriguingly, time-lapse imaging of Wnt/CWN-1::mNG showed primarily anterograde movement of Wnt particles in both the anteriorly and posteriorly directed CAN processes (Video S2), along the length of the animal, which suggests these cells may have the ability to present Wnt ligands to much of the body. The CANs are an enigmatic cell type with long axon-like processes that resemble neurons but lack synapses [5]. *C. elegans* has one CAN cell on each side, each with a cell body located near the vulva and two axon-like, microtubule-based structures [6] that reach nearly the entire length of the body, to the head and tail. CANs do not have known roles in neurotransmission, but instead have essential functions in development, growth, and survival [7, 8], which is consistent with a function for CAN cell architecture in mediating long-range signaling. To test the extent to which Wnt transport in neurons may generally resemble Wnt/CWN-1 transport in CAN cells, we next examined Wnt/EGL-20 dynamics in *egl-20* expressing neurons with axons that projected along the ventral nerve cord. We observed that Wnt/EGL-20::mNG accumulated on cells adjacent to these axons (Fig. 1d), but Wnt/EGL-20::mNG particles in *egl-20*-expressing axons themselves were rare and non-mobile, which suggested the mode of transport differs from Wnt/CWN-1 transport in CAN. To test whether these differences in Wnt transport are due to cell type or ligand, we expressed EGL-20::mNG in CAN cells using a short fragment of the *cwn-1* promoter (Fig. 1e). Transgenically expressed *Pcwn-1*^*(- 272)*^>EGL-20::mNG localized to mobile punctae in CAN cell extensions similar to endogenous Wnt/CWN-1::mNG. This observation suggests that the transport processes for these two Wnt proteins in their endogenous contexts likely depend on cell type-specific mechanisms rather than intrinsic properties of the different ligands.

**Figure 1.**
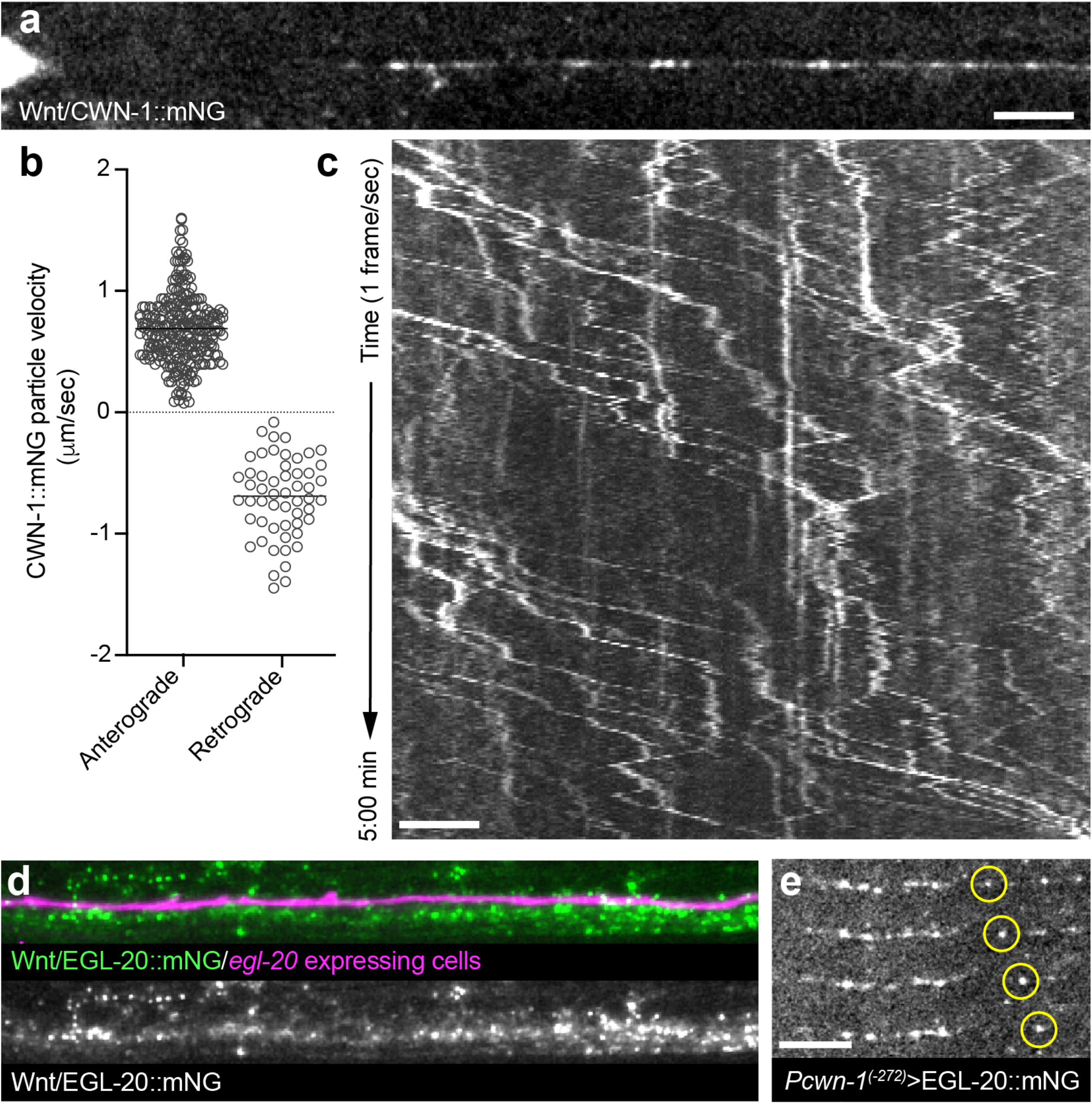
Wnt trafficking in CAN highlights cell type-specific modes of long-distance Wnt transport *in vivo*. Live imaging of endogenously tagged Wnt/CWN-1::mNG and Wnt/EGL-20::mNG reveals an unexpected mode of Wnt transport in CAN cells and demonstrates cell type-specific Wnt transport processes. (**a**) CWN-1::mNG localized to prominent particles in a CAN axon. Image corresponds to Video S1 and the first line of the kymograph in (c); (**b**) quantification of CWN-1::mNG particle velocities in CAN axons based on multiple tracks from 11 animals; (**c**) kymograph depicting CWN-1::mNG dynamics within a CAN axon based on time-lapse imaging (see Video S1). Images were acquired every one second for five minutes; (**d**) localization of EGL-20::mNG near the ventral nerve cord in a living animal. The axon of an *egl-20*-expressing tail neuron is highlighted by *Pegl-20*>2x mKate2::PH; (**e**) time-lapse imaging showing EGL-20::mNG is transported as mobile particles when transgenically expressed in CAN. Image shows four frames taken at one second intervals with a mobile EGL-20::mNG particle circled. All scale bars = 5 μm

Our finding that a Wnt is directionally trafficked over long distances in axon-like structures, to our knowledge, represents a novel process for long distance Wnt transport *in vivo*. Despite widespread potential for Wnt signaling from neurons to other cell types in many animals, there are relatively limited data on developmental and homeostatic functions for Wnts produced by differentiated neurons [9, 10] or neuron-like cells, which may be a fruitful avenue for future research. The Wnt transport mechanisms we discovered likely differ from transport mechanisms in cytonemes given the cytoskeletal differences between dynamic, actin-based cytonemes and stable, microtubule-based structures such as axons, dendrites, and CAN cell processes [6]. Our comparison of CWN-1 and EGL-20 dynamics, along with EGL-20 misexpression in CAN cells, also provides direct evidence that Wnt proteins utilize distinct transport modes *in vivo* depending on the cells that they are expressed in. These findings highlight the diversity of mechanisms involved in long-range Wnt signaling in living animals and the importance of live imaging endogenous proteins to reveal these mechanisms. We propose that neuronal cell architectures may provide easily repurposed routes for long-distance Wnt transport that complement other dispersal processes such as diffusion or cytonemes. In particular, Wnt delivery by neurons and neuron-like cells could enhance local signaling precision within organismal scale Wnt gradients generated by diffusion.

## Acknowledgments

We thank the members of the Pani and Goldstein labs for helpful discussions. Some strains were provided by the CGC, which is funded by NIH Office of Research Infrastructure Programs P40 OD010440.

## Author contributions

AMP: conceptualization, investigation, formal analysis, writing – original draft, writing – review and editing, funding acquisition; MF: investigation, formal analysis; BG: funding acquisition, resources, writing – review and editing.

## Supplementary Materials

### Videos

**Video S1**

Time lapse imaging of endogenously tagged Wnt/CWN-1::mNG dynamics illustrating directional transport along an axon-like CAN structure. Images were acquired every one second for five minutes using a spinning disk confocal microscope.

Available at: https://figshare.com/articles/media/Pani_Favichia_Goldstein_2023_-_Video_S1/22732760

**Video S2**

Time lapse imaging of endogenously tagged Wnt/CWN-1::mNG demonstrating anterograde transport in both the anteriorly (left) and posteriorly (right) directed CAN processes. Images were acquired every one second for 30 seconds using a spinning disk confocal microscope.

Available at: https://figshare.com/articles/media/Pani_Favichia_Goldstein_2023_-_Video_S2/22732829

## Materials and methods

### *C. elegans* maintenance, gene tagging and transgenesis

*Caenorhabditis elegans* were grown on Nematode Growth Media (NGM) plates, fed *E. coli* (OP50 strain), and cultured at 20°C for experiments. To generate *cwn-1(cp209[cwn-1::mNG::3xFlag])*, we inserted mNG::3xFlag at the C-terminus of *cwn-1* using Cas9-triggered homologous recombination with a self-excising selection cassette as described previously [11]. Homology arms were cloned into pDD268 [11] (Addgene #132523) using NEBuilder Hifi DNA assembly Master Mix (New England Biolabs). The guide RNA targeting sequence 5’- TGGTCGTGGATACGAGAAGA-3’ was inserted into the *Peft-3*>Cas9 + sgRNA plasmid pDD162 [12] (Addgene #47549) using a Q5 site-directed mutagenesis (New England Biolabs). CWN-1::mNG::3xFlag knock-in animals were phenotypically indistinguishable from wild-type. Endogenously tagged EGL-20::mNG::3xFlag animals are also indistinguishable from wild-type [2]. The *Pcwn-1*^*(-272)*^ promoter used for CAN expression in pAP181 and pAP209 consisted of 272 base-pairs of genomic sequence upstream of the *cwn-1* translation start site.

Transgene constructs were assembled in the pAP088 backbone [2] for single-copy insertions near the ttTi4348 site on Chr I using NEBuilder Hifi DNA assembly Master Mix (New England Biolabs). The *egl-20* coding sequence in pAP209 was codon-optimized using the *C. elegans* codon adapter tool [13] and synthesized as a gBlock DNA fragment (Integrated DNA Technologies).

### Live imaging and analyses

Larval animals were immobilized for imaging using 0.1 mmol/L levamisole in M9 buffer and mounted on 3% (wt/vol) agarose pads. Animals were imaged within 30 minutes of mounting, and data shown are representative of at least 10 animals imaged on at least two occasions. Numerous animals were mounted on each slide, and individuals were chosen for imaging based on developmental stage, orientation on the slide, and health. Imaging systems utilized included: (1) Nikon Ti-E stand with CSU-X1 spinning disk confocal (Yokogawa), 447, 514, and 561 nm solid state lasers, ImageEM EMCCD camera (Hamamatsu) and 100x 1.49 NA objective; (2) Nikon Ti-2 stand with CSU-X1 spinning disk confocal (Yokogawa), 445, 514, and 561 nm solid state lasers, ORCA Fusion BT sCMOS camera (Hamamatsu) and 100x 1.49 NA objective; and (3) Nikon Ti-2 stand with CSU-W1 SoRa spinning disk confocal (Yokogawa), 445, 514, and 594 nm solid state lasers, ORCA Fusion sCMOS camera (Hamamatsu) and 60x 1.27 or 100x 1.49 NA objectives. Images were acquired using MetaMorph (Molecular Devices) or NIS Elements (Nikon) software with varying camera exposure times and laser intensities depending on the strain and imaging system. Kymographs were created using the KymoResliceWide plugin (https://github.com/UU-cellbiology/KymoResliceWide) for FIJI [14]. Particle velocities for CWN-1::mNG transport in CAN were calculated from kymographs by manual tracing of frame-to-frame paths for multiple tracks in 11 animals. Figures were prepared using FIJI [14], Prism 9.5 (GraphPad), and Adobe Illustrator 24.1 (Adobe).

### *C. elegans* strains and reagents

| Reagent type | Description | Identifier | Source | Additional information |
| --- | --- | --- | --- | --- |
| *C. elegans* strain | lfpls4[Pcwn-1(-272)>flp::SL2::2x mTurq2uoise::PH::3xHA::tbb-2 3'UTR LoxN] I; cwn-1(cp209[cwn-1::mNG-C1^3xFlag]) II | LP851 | This study |  |
| *C. elegans* strain | cpls89[Pwrt-2>2x mTurquoise2::PH::tbb-2 3'UTR LoxN] I; cpls85[Pegl20>2x mKate2::PH::3xHA::tbb-2 3'UTR loxN] II; egl-20(cp217[egl-20::mNG^3xFlag]) IV | LP515 | Pani and Goldstein, 2018, Ref. 2 |  |
| *C. elegans* strain | lfjEx1 (Pcwn-1(-272)>egl-20::mNG::SL2::2x mTurquoise2::PH::3xHA::tbb-2 3'UTR | APL185 | This study |  |
| Recombinant DNA reagent | Peft-3>Cas9::tbb-2 3'UTR + PU6>empty sgRNA | pDD162 | Dickinson, *et al*, 2013, Ref. 12 | Plasmid for Cas9 + sgRNA cloning |
| Recombinant DNA reagent | ccdB mNG^SEC^3xFlag ccdB | pDD268 | Dickinson, *et al*, 2015, Ref. 11 | Plasmid for cloning mNG::3xFlag homologous repair templates |
| Recombinant DNA reagent | cwn-1::mNG^SEC^3xFlag | pAP079 | This study | Homologous repair template to generate endogenous cwn-1::mNG::3xFlag knock-in |
| Recombinant DNA reagent | Peft-3>Cas9::tbb-2 3'UTR + PU6>cwn-1C sgRNA | pAP080 | This study | Cas9 + sgRNA plasmid targeting c-terminus of *cwn-1* |
| Recombinant DNA reagent | Peft-3>Cas9::tbb-2 3'UTR + PU6>ttTi4348 sgRNA | pAP082 | Pani and Goldstein, 2018, Ref. 2 | Cas9 + sgRNA plasmid targeting genomic site near ttTi4348 on Chr I |
| Recombinant DNA reagent | empty promoter>2x mTurquoise2::PH::3xHA::tbb-2 3'UTR loxN SEC loxN ttTi4348 | pAP088 | Pani and Goldstein, 2018, Ref. 2 | Plasmid for plasma membrane marker insertions near ttTi4348 |
| Recombinant DNA reagent | Pcwn-1(-272)>flp::SL2::2x mTurquoise2::PH::3xHA::tbb-2 3'UTR loxN SEC loxN ttTi4348 | pAP181 | This study | Plasmid expressing mTurquoise2::PH membrane marker and flp in CAN cells |
| Recombinant DNA reagent | Pcwn-1(-272)>flp::SL2::2x mTurquoise2::PH::3xHA::tbb-2 3'UTR loxN SEC loxN ttTi4348 | pAP209 | This study | Plasmid expressing egl-20::mNG and mTurquoise2::PH membrane marker in CAN cells |

